# Behavior of Neural Cells Post Manufacturing and After Prolonged Encapsulation within Conductive Graphene-Laden Alginate Microfibers

**DOI:** 10.1101/2021.04.04.438421

**Authors:** Marilyn C. McNamara, Amir Ehsan Niaraki Asli, Rajeendra L. Pemathilaka, Alex H. Wrede, Reza Montazami, Nicole N. Hashemi

## Abstract

Engineering conductive 3D cell scaffoldings offer unique advantages towards the creation of physiologically relevant platforms with integrated real-time sensing capabilities. Toward this goal, rat dopaminergic neural cells were encapsulated into graphene-laden alginate microfibers using a microfluidic fiber fabrication approach, which is unmatched for creating continuous, highly tunable microfibers. Incorporating graphene increases the conductivity of the alginate microfibers 148%, creating a similar conductivity to native brain tissue. Graphene leads to an increase in the cross-sectional sizes and porosities of the fibers, while reducing the roughness of the fiber surface. The cell encapsulation procedure has an efficiency rate of 50%, and of those cells, approximately 30% remain for the entire 6-day observation period. To understand how encapsulation effects cell genetics, the genes IL-1β, TH, TNF-α, and TUBB-3 are analyzed, both after manufacturing and after encapsulation for six days. The manufacturing process and combination with alginate leads to an upregulation of TH, and the introduction of graphene further increases its levels; however, the inverse trend is true of TUBB-3. Long-term encapsulation shows continued upregulation of TH and of TNF-α, and six-day exposure to graphene leads to the upregulation of TUBB-3 and IL-1β, which indicates increased inflammation.

## 1. Introduction

While three-dimensional (3D) cell cultures continue to grow in complexity and physiological relevance, more work must be done to reach the full potential of a real-time cell sensing system that is able to match the macro- and microenvironments of target tissues. One-dimensional (1D) and two-dimensional (2D) real-time sensors have been reliably created utilizing micro- and nano-electrodes, or planar electrodes, respectively.^[1, 2]^ This work furthers the cause by using biocompatible, graphene-laden microfibers as cellular constructs, which can be used in conjunction with 3D micro-electrode arrays (MEAs) for a highly complex real-time sensing system to analyze electrical cell-to-cell communication that occurs within the brain. Additionally, this study works towards the important task of identifying genetic changes caused by manufacturing, and contrasting this against the effects of long-term encapsulation in four genes that are important to neural health, such as tyrosine hydroxylase (TH), tubulin beta 3 class 3 (TUBB-3), interleukin 1 beta (IL-1β), and tumor necrosis factor alfa (TNF-α).

Hydrogels, with their high water content and the ease of diffusion across their borders, are ideal candidates for applications wherein the spatiotemporal properties of the cells must be controlled for long-term observation.^[3, 4–7]^ In particular, microfibers are well-suited for this purpose, as their higher surface-to-volume ratio expedites the diffusion of nutrients and waste across the cell border, while allowing for highly complex and specific scaffold geometries.^[4, 7–10]^ Cell-laden microfibers can be created in a number of different ways, including wetspinning/extrusion;^[11]^ however, microfluidics provides unmatched control over the size, shape, and degredation rates of the resulting microfibers, while still allowing for all potential cell-safe gelation methods.^[8, 12]^ In this way, a cell suspension might be mixed with a pre-gel solution before polymerization or gelation, thereby resulting in a cell-laden microfiber with highly controlled mechanical properties.

In this work, alginate was chosen for its cell compatibility, but its low conductivity needed to be addressed. There are a number of synthetic or carbon-based materials that can enhance the naturally low conductivity of alginate; these materials include synthetic polymers such as PEDOT:PSS,^[13]^ silver nanowires,^[14]^ graphite,^[15, 16]^ graphene oxide,^[17, 18]^ and pure graphene.^[9, 18, 19]^ While pure graphene is favorable due to its extraordinary electrical, optical, biomedical, and mechanical properties, incorporating it into a stable aqueous pre-polymer solution for use with cell encapsulation is difficult, as the creation of graphene solutions typically requires the use of toxic or unstable chemicals, such as dimethylformamide (DMF) or N-methylpyyrolidone (NMP).^[1, 20]^ Fortunately, bovine serum albumin (BSA), an edible protein, can be used to stabilize aqueous graphene solutions after exposing graphite to mechanical shear stress through liquid phase exfoliation (LPF).^[1, 9, 21]^ This mitigated the need for cytotoxic chemicals to stabilize a graphene dispersion. In this study, shear stress was created through the use of a kitchen blender in a cost-effective and scalable method.^[1, 22, 23]^

Any cell-laden 3D sensing system must be physiologically and genetically relevant to the target tissue in order to avoid falsely skewing results. In order to determine the genetic effects of encapsulating cells within alginate and graphene-alginate microfibers, real time reverse transcription quantitative polymerase chain reactions (RT-qPCR) was carried out on rat dopaminergic neural cells (N27s), which were encapsulated into alginate or graphene-alginate microfibers for up to six days. Genes of interest for understanding the health and behavior of neural cells include TH, TUBB-3, IL-1β, and TNF-α. TH is a rate-limiting enzyme in dopamine synthesis, is a crucial key to understand and monitor the health of dopaminergic N27 cells,^[24]^ while TUBB-3 is responsible for forming the cellular cytoskeleton, aiding in cell migration and organellar transport, and neurogenesis.^[25]^ To better understand the effects of manufacturing on the genetic makeup of encapsulated cells, RT-qPCR was used both after manufacturing and after prolonged encapsulation for TH and TUBB-3.

Similarly, IL-1β and TNF-α were observed after six days of encapsulation to identify potential sources of inflammation within encapsulated N27 cells. Neural cells react to negative conditions by increasing microglia activation to create an inflammatory response; as such, studying the expression of IL-1β and TNF-α are excellent ways to understand the health of a neural tissue culture.^[26]^ While IL-1β is an important mediator of the inflammatory response, TNF-α is a well-characterized cytokine involved with inflammation, and is specifically involved with the acute phase reaction; widespread upregulation of TNF-α can lead to systemic inflammatory disorders such as rheumatoid arthritis and Crohn’s disease.^[27]^ Studying both genes after prolonged encapsulation allows for a thorough understanding of the inflammatory response of N27 cells encapsulated into alginate or graphene-alginate microfibers. Past work from Song et al. (2014) has shown that microglia (BV2) cells cultured onto 3D graphene foams showed an increase in IL-1β and TNF-α levels in the absence of lipopolysaccharides.^[26]^

This work features a novel method for the fabrication of graphene- and cell-laden microfibers using a microfluidic fiber fabrication method. Introduction of graphene into the alginate polymer changed the size, porosity, and surface topographies of the resulting microfibers, thereby potentially adjusting the diffusion potential for waste and nutrients to exit and enter the fiber boundary, and potentially affecting cellular viability and genetic expression.^[8]^ Importantly, viability and genetic expression were investigated both shortly after manufacturing and after prolonged encapsulation, to better understand the implications of the manufacturing process compared to the encapsulation of the cells. Elucidation of this facet of cellular encapsulation is crucial to furthering the technology of 3D culturing, both with and without conductive elements.

## 2. Results

### 2.1 Microfluidic Microfiber Fabrication Overview

Microfluidic fiber fabrication relies upon the use of a microfluidic chip, variations of which have commonly been used for biomedical applications.^[28]^ Changing the geometry and number of inlets can drastically affect the shape and size of the resulting fibers.^[8, 29, 30, 31]^ During fiber fabrication, both core (pre-gel) and sheath (gellator) solutions are used; the core solution will be crosslinked into the resulting microfiber, while the sheath solution helps guide the core solution through the channel, further shaping it and preventing clogging.^[6, 30, 32, 33]^ The viscosities and flow rates used can also affect the shape, size, topography, and mechanical properties of the resulting fibers.^[32]^ In this case, the sheath solution also contains ionic crosslinkers that cause the gellation of the core fluid before it leaves the microfluidic chip; upon exiting the chip, the newly formed microfiber will be introduced into a collection bath that will further strengthen it and allow for easy gathering.

For this work, the core solution was created with alginate, a commonly used biopolymer derived from the cell walls of brown algea.^[7, 32, 34]^ Alginate features favorable biocompatibility parameters such as cell-safe gelation conditions, high permeability, neutral pH, and low antigenicity.^[16]^ Its structure of M- and G-blocks can be ionically crosslinked in the presence of divalant cations, such as Ca^2+^, Ba^2+^, Sr^2+^, and others; however, calcium crosslinkers are commonly used in biomedical applications because they are readily available and cost effective, and because calcium alginate hydrogels exhibit good swelling properties, which is important to maintain cell viability.^[5, 6, 9, 35]^

### 2.2 Viscosities of Alginate and Graphene-Alginate Solutions

The viscosities of cell-free pre-polymer solutions were measured using a Cannon-Fenske viscometer, as previously described.^[32]^ The addition of graphene into the 3.5% alginate core solution significantly increased the viscosity of the fluid, which can be seen in **Figure 1** (A). This can be attributed to the attachment of alginate chains onto the graphene flakes.^[20]^ A thicker, more viscous core solution is more resistant to shear force from the sheath fluid within the microfluidic device, which can significantly affect the properties of the resulting fibers.

**Figure 1.**
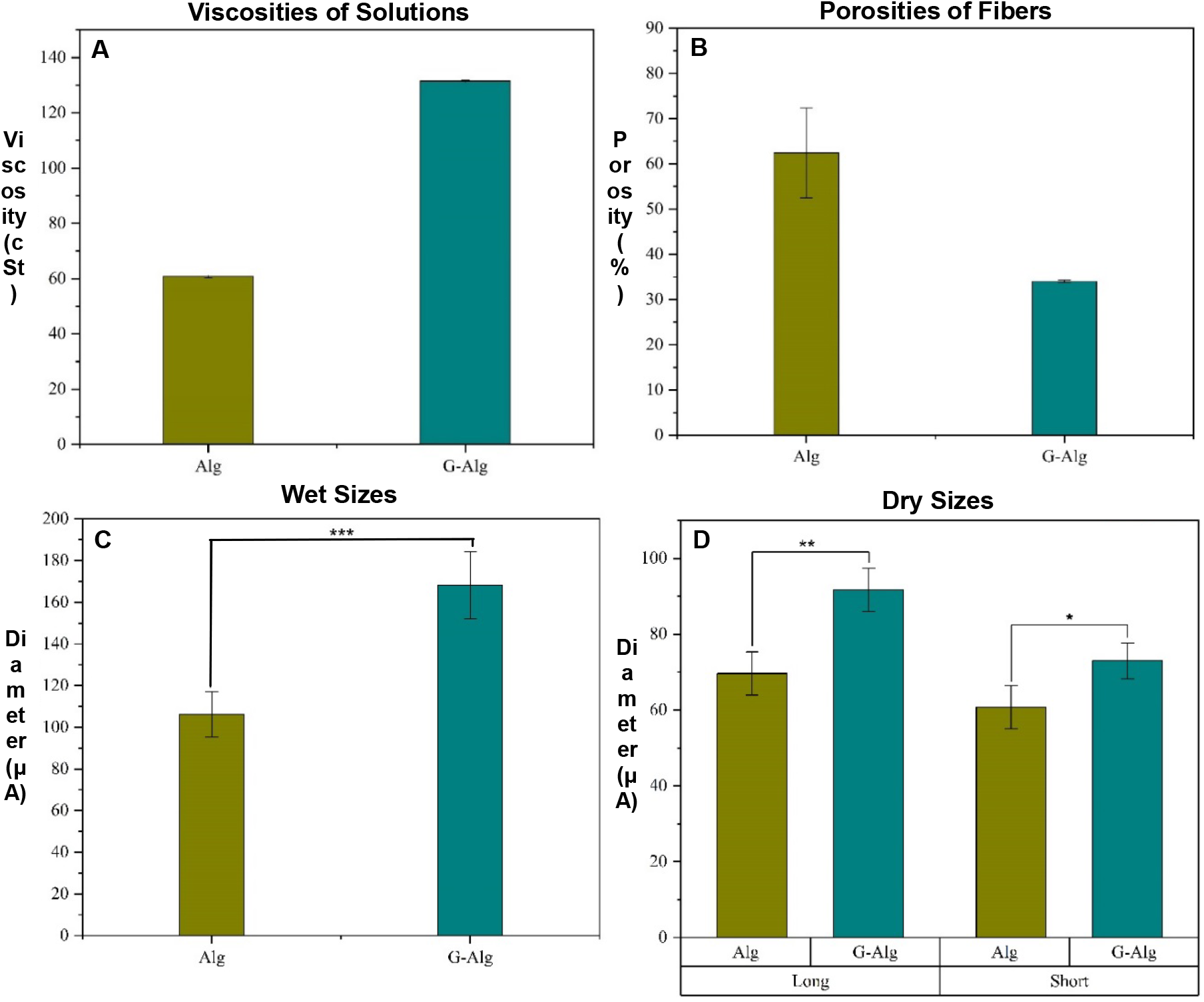
(**A**) Viscosities of 3.5% alginate (Alg) and 3.5% alginate in graphene (G-Alg) solutions, as measured by a Cannon-Fenske viscometer. Error bars represent ± 1 standard deviation. (**B**) Porosities of alginate (Alg) and graphene-alginate (G-Alg) microfibers. Error bars represent ± 1 standard deviation. (n ≥ 3). (**C-D**) Sizes of wet (**C**) and dry (**D**) microfluidically-created alginate and graphene-alginate microfibers. Dry fibers have oval-shaped cross-sectional areas, and therefore have both a long and a short dimension. Error bars represent ± 1 standard deviation. (n ≥ 10, \**p* ≤ 0.05; ** *p* ≤ 0.01; *** *p* ≤ 0.001).

### 2.3 Sizes, Surface Topographies, and Porosities of the Microfibers

The porosities of alginate microfibers were significantly higher than those of graphene-alginate microfibers (Figure 1 (B)). This might be attributed to the rougher surface topography, which would indicate a higher surface area and therefore increased opportunities for diffusion across the fiber boundary. Likewise, it is possible that the addition of graphene into the polymer matrix affected the amount of water the fibers could absorb due to increased interactions between the alginate and graphene particles.^[20]^ Since the graphene solutions are significantly more viscous than their alginate counterparts (Figure 1 (A)), it follows that the within the microfluidic device, graphene solutions would be more resistant to deformation by shear force exerted from the sheath fluid.^[32]^ Therefore, the sizes of both wet and dry graphene-alginate microfibers are larger than the corresponding alginate microfibers (Figure 1 (C-D)).

Likewise, an examination of the surface roughness (R_a_) of the surface of graphene-alginate and alginate microfibers shows that the shear force has a lesser effect on the topographies of graphene-alginate microfibers as opposed to alginate microfibers (**Figure 2**). The surface roughness measurement R_a_ is defined as the average of the distance from the mean vertical value of the topography across a given sample.^[32]^ Graphene-alginate microfibers were significantly smoother across the surface of the fiber, but both fibers exhibited the same R_a_ along the length of the fibers, with both samples showing on average less than 4 μm of distance from the mean vertical value. This speaks to the consistency of the continuous fiber fabrication achieved by the microfluidic method.

**Figure 2.**
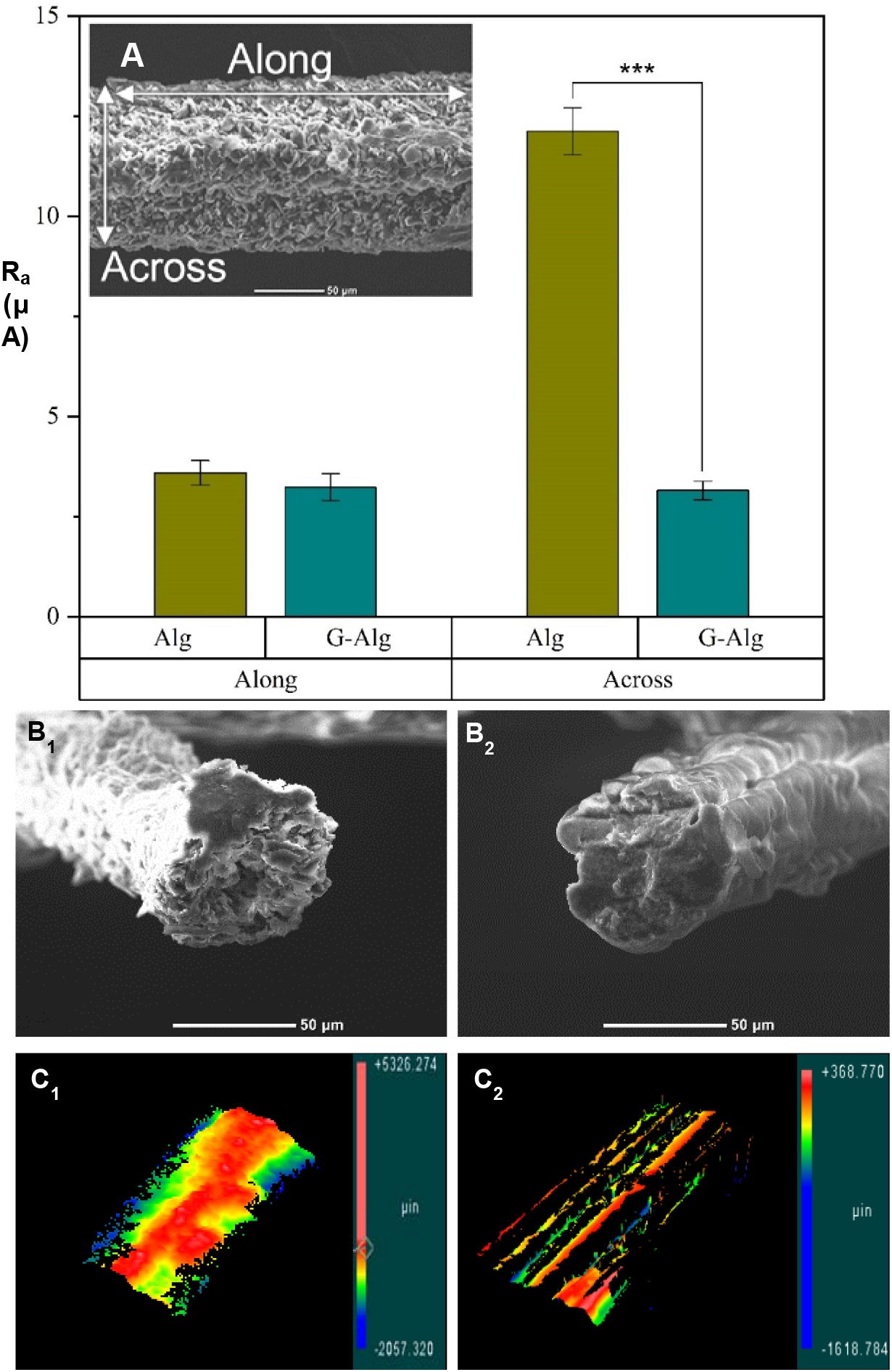
(**A**) Quantifications of the surface roughness (R_a_) of alginate (Alg) and graphene-alginate (G-Alg) microfibers, both along and across the fibers. Along and across are demonstrated on an SEM image of a graphene-alginate microfiber. Error bars represent ± 1 standard deviation. (n ≥ 7, ***p ≤ 0.001). (**B**) SEM images of the cross-sections of graphene-alginate (1) and alginate (2) microfibers. (**C**) 3D topologies of the surfaces of dry graphene-alginate (1) and alginate (2) microfibers, taken with a profilometer.

Overall, the graphene-alginate fibers had a lower porosity and surface roughness and a larger size than their pure alginate counterparts; combined, these parameters affect the amount of fluids that are able to enter or exit the fiber boundaries. Graphene-alginate fibers might have a lower amount of nutrients diffusing from the maintenance media into the encapsulated cells, and waste might not be able to exit the fibers as easily, thereby potentially creating a more difficult living situation for cells encapsulated into graphene-alginate microfibers. This might affect the viability and genetic expressions of cells encapsulated into graphene-alginate microfibers when compared against those in alginate fibers.

### 2.4 Conductivity and Cyclic Voltammetry

In order to better understand the electrical properties of the microfibers, multiple cyclic voltammetry tests were carried out upon using a potentiostat. The results show little to no change over the course of ten cycles, and the slight difference between the charge and discharge portions indicate that all samples had low capacitance. Similarly, most samples exhibit very weak oxidation/reduction peaks, particularly within the dry samples.

Introducing graphene into the alginate microfibers did not drastically affect the conductivity of dry fibers, which were mounted onto polystyrene slips with carbon tape and silver paste (**Figure 3** (A-C)). However, when the fibers were soaked in water overnight and measured again, there was a significant change in the conductivities of the wet samples, with wet graphene-alginate microfibers showing a significantly higher conductivity than pure alginate microfibers. This is due to the interaction between water and graphene; when present, water can amplify ion exchange within the polymer.

**Figure 3.**
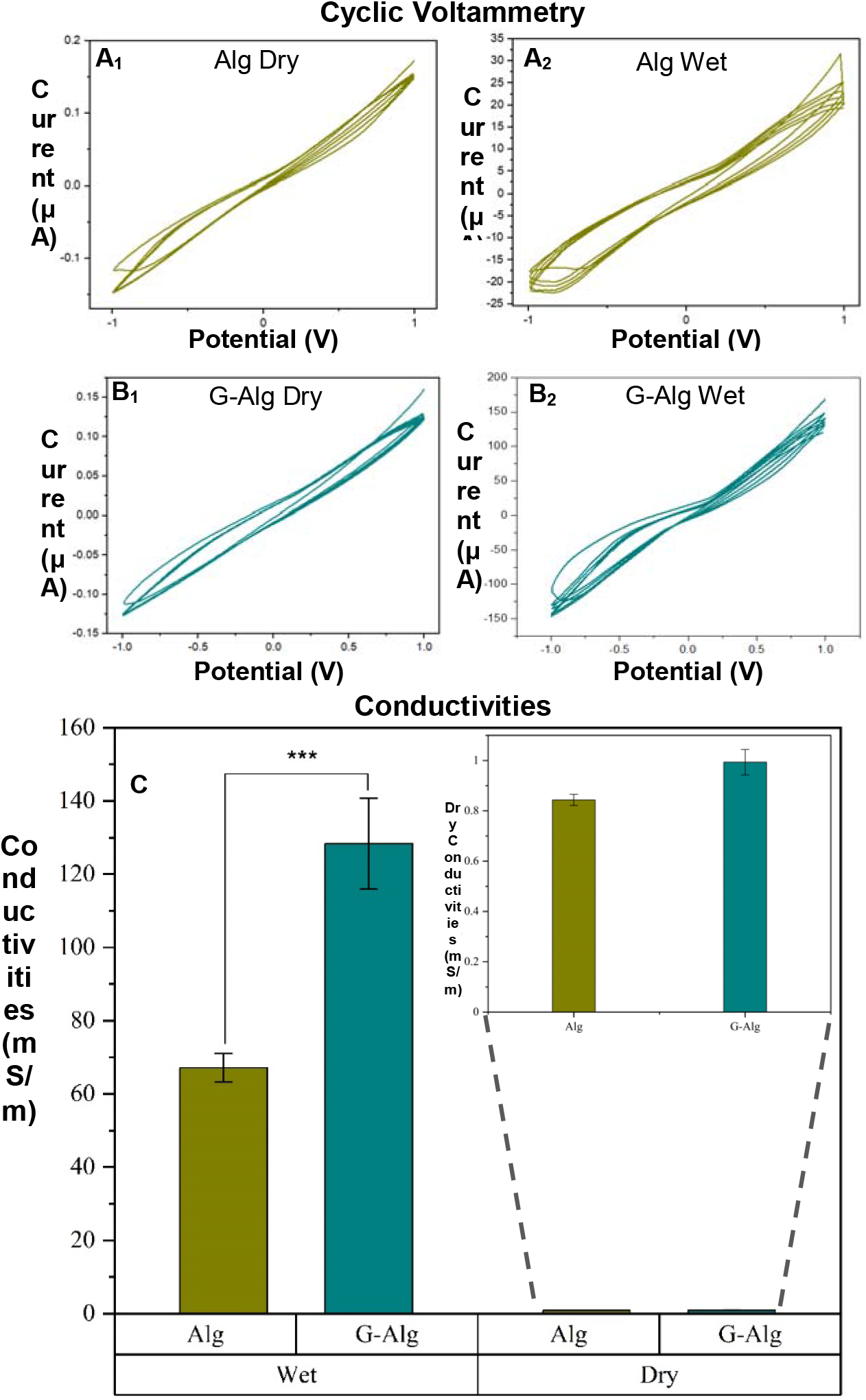
(**A-B**) Cyclic voltammetry of alginate (A) and graphene-alginate (B) microfibers, collected by a potentiostat from −1 V to 1 V at a rate of 0.1 V s-1. Samples were tested both dry (A_1_-B_1_) and wet (A_2_-B_2_). (**C**) Conductivities of both wet and dry alginate and graphene-alginate microfibers. Fibers were mounted onto nonconductive polysterine with carbon tape, silver paste, and copper leads so that cyclic voltammetry could be carried out with a potentiostat. Error bars represent ± 1 standard deviation. (n ≥ 5, *** *p* ≤ 0.001).

The values presented in this work are favorable when compared with other studies in which cells are encapsulated within conductive hydrogels. In each instance, researchers attempted to match the conductivities of the native tissues. For instance, Dong et al. (2018) encapsulated cardiac cells into chitosan-graft-aniline tetramer (CS-AT) with dibenzaldehyde-terminated poly(ethylene glycol) (PEG-DA) to create conductive, self-healing, cell-laden hydrogels with a conductivity of 200 mS m^−1^.^[36]^ Similarly, Sirivisoot et al. (2014) achieved a conductivity of 10 mS m^−1^ when encapsulating PC12 cells for the treatment of nerve injuries.^[37]^

For native brain tissue, weighted average means from a current meta-analysis indicate that brain tissues have conductivities in the range of 220 mS m^−1^ to 470 mS m^−1^, for white and gray matter, respectively.^[38]^ Therefore, the addition of graphene into alginate microfibers brought the wet conductivity of the resulting hydrogel close to the range of the native target values, increasing the conductivity from 67.1 ± 3.9 mS m^−1^ to a final value of 128.4 ± 12.4 mS m^−1^ for an increase of 148.4% (Figure 3 (C)).

### 2.5 Encapsulation of N27 Cells within Alginate and Graphene-Alginate Microfibers

Cells encapsulated in microfibers were observed in the alginate-based solution, immediately after fabrication, and on days one, four, and six. All values were normalized against the weight of alginate solutions or fibers to provide an easy way to compare across all samples. Cell suspensions were mixed with alginate solutions to a final concentration of 2 x 10^6^ cells mL^−1^; however, one mL of 3.5% alginate weighed in at 1.1 grams, so the initial cell concentration was 1.82 x 10^6^ cells g^−1^, as seen in **Figure 4**.

**Figure 4.**
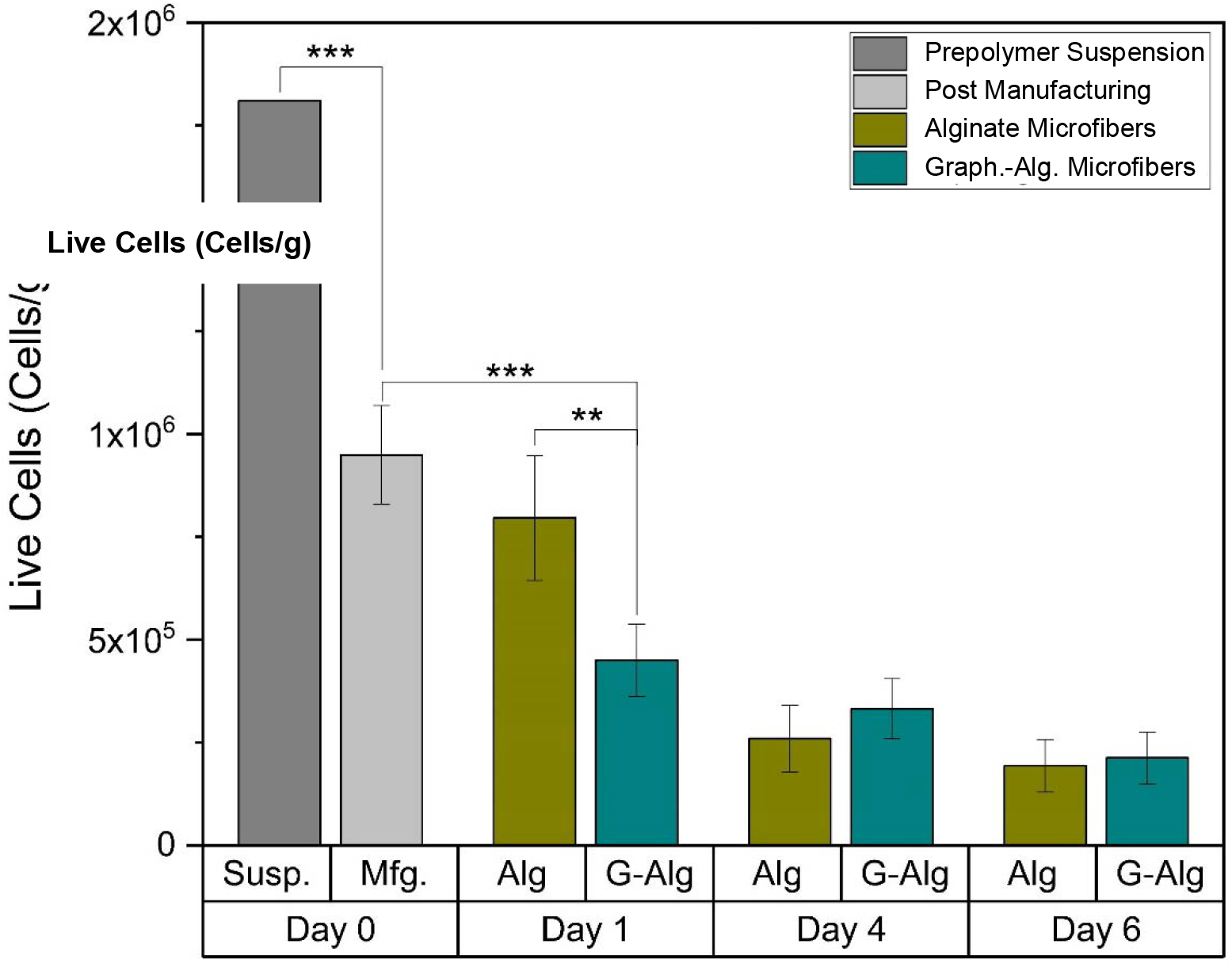
Numbers of live cells within relevant experimental samples of alginate solutions and alginate or graphene-alginate microfibers. The numbers of live cells were calculated using a Trypan blue exclusion. Error bars represent ± 1 standard deviation. (n ≥ 5, ** *p* ≤ 0.01; *** *p* ≤ 0.001).

Counting the cells in fibers immediately after fabrication showed a concentration of approximately 1 x 10^6^ cells g^−1^, which indicates that cells were roughly half of the cells in the initial solution did not get encapsulated within the microfibers. This loss can be attributed to multiple factors; firstly, cells in the prepolymer solution settle down when in the syringe, and therefore some will be lost before they can enter the microfluidic device. Additionally, some cells will not survive the manufacturing process; if they are too near the fluid-fluid boundary, they may experience adverse conditions, such as shear force.

Most cells survived the first day when encapsulated into pure alginate microfibers. However, there was some drop in the number of viable cells within alginate-graphene microfibers; this might be caused by the aforementioned bonds between alginate and graphene, which increased the viscosity of the prepolymer solutions, which led to the increased size and decreased porosity and surface roughness of graphene-alginate microfibers, when compared against pure alginate microfibers. These factors could inhibit diffusion into and out of the fiber boundary, causing difficulties getting nutrients into and waste out of the fiber, thereby leading to cell death. However, on days four and six, the difference between live cells within alginate and graphene-alginate microfibers is insignificant, likely due to some natural degradation fibers experience within the media allowing for better nutrient permeation to the cells. Days four and six also saw a drop in the number of live cells when compared with day one. While some cells might die within the fibers, it is also possible for them to migrate beyond the fiber borders, where they can be observed attached to the bottom of the well plates.

### 2.6 Genetic Expressions of N27 Cells Post Manufacturing (Day One)

The goal of this study was to piece apart the genetic effects of the introducing N27 cells into alginate or graphene-alginate solutions, the actual microfluidic fiber fabrication process, and long-term (six-day) encapsulation within alginate or graphene-alginate hydrogels. The novelty of this work is due to the focus on the actual encapsulation process, and how it affects the genetic expressions of genes such as TH or TUBB-3, which are important markers of nerual health.

In order to understand the effects of manufacturing on genetic expression, samples were tested one day after encapsulation to allow for the appropriate amount of time for genetic changes to take effect after an event which might affect the genetic makeup of the cells. An ‘Alginate Control’ sample was gathered to understand what genetic changes could be attributed to the effects of alginate or the temperature changes the cells experience through encapsulation without going through the microfluidic device or coming in contact with CaCl_2_·2H_2_O. In contrast, the day one samples experienced the full encapsulation procedure, and were cultured at maintenance conditions for 24 hours. In this way, the individual pieces of manufacturing are understood in relation to the genetic expression of the cells.

Two genes were studied to understand how the manufacturing process affected the genetic expression of N27 cells. TH is a key dopaminergic neuron marker^[39]^ that helps with the production of dopamine, and is therefore crucial to the health of the nervous system, and is incredibly important in the development of neurodegenerative diseases, like Parkinson’s disease.^[40]^ Increased amounts of TH can cause oxidative stress that can create a toxic environment for cells.^[41]^ Additionally, TUBB-3, a gene which encodes for a class III member of the beta tubulin protein family, is related to neurogenesis, as well as axonal growth and maintenance.^[25]^ Abnormally high upregulation of TUBB-3 in tissues might be an indication of cancer.^[42]^ It can be crucial in recovery from traumatic brain injuries. The genetic effects of the manufacturing process on the genes TH and TUBB-3 can be observed in **Figure 5.**

**Figure 5.**
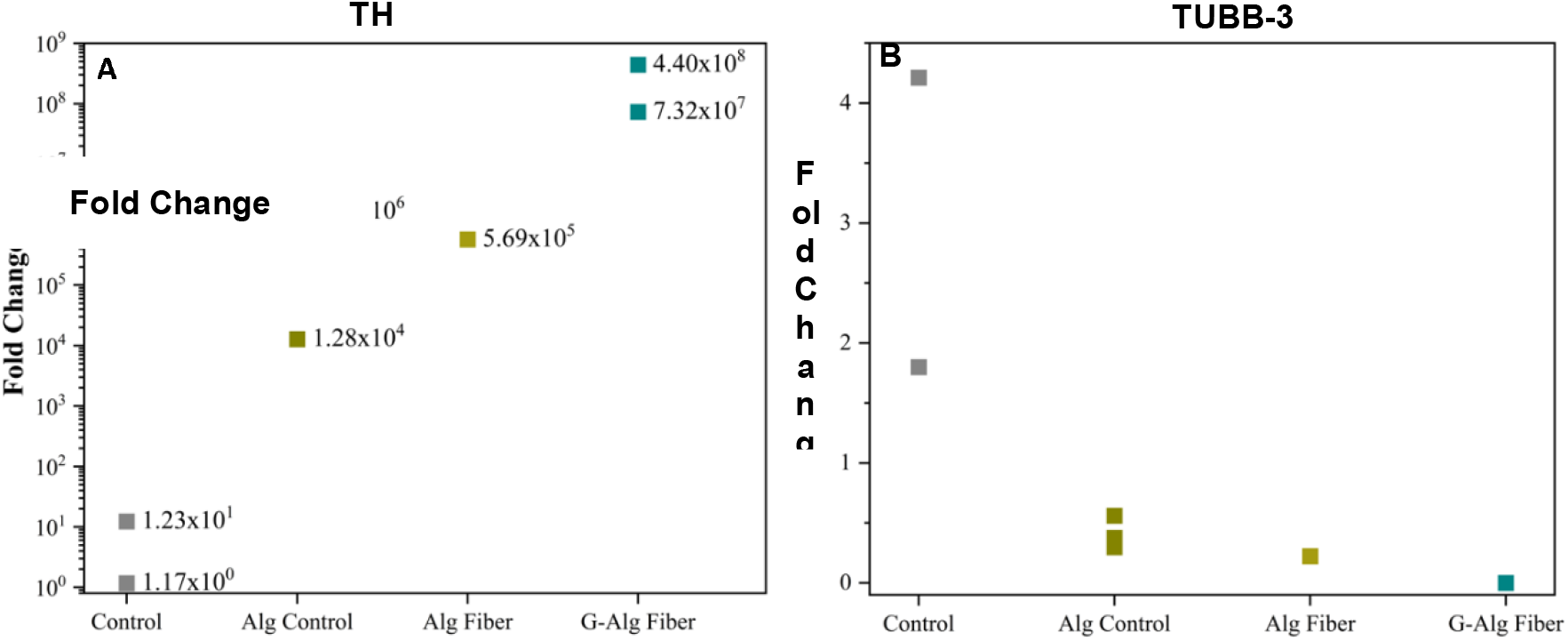
RT-qPCR analysis of the expression levels of TH **(A)** and TUBB-3 **(B)** within N27 cells 24 hours after seeding (Control, Alg Control) or encapsulation (Alg Fiber, G-Alg Fiber). (n≥1).

TH was greatly upregulated within the alginate control samples, indicating that alginate itself and the temperature changes the cells experience during an encapsulation experiment increased the expression of TH within the N27 cells when compared against controls (Figure 5 (A)). However, levels within alginate microfibers after 24 hours were not greatly different than in the alginate controls, indicating that the rest of the manufacturing procedure did not greatly affect the TH expression levels. N27 cells in graphene-alginate microfibers did express higher levels of TH than those in pure alginate microfibers.

Therefore, when compared against normal N27 cells, cells that were in contact with alginate solutions or had just undergone microfluidic encapsulation into alginate hydrogels show increased TH expression and dopaminergic activity, and cells that were freshly encapsulated into graphene-alginate microfibers show a further increase of TH expression, indicating that cells in contact with alginate and graphene were distress and were potentially exhibiting higher levels of dopamine release, which could in turn lead to the increased cell death that was observed in day one graphene-alginate samples. Similarly, Tasnim et al. (2018) have shown that culturing mesenchymal stem cells (MSCs) onto graphene foams leads to an increase of TH expression, indicating that the graphene encourages seeded MSCs to differentiate into dopamine-producing dopaminergic neurons.^[43]^

Conversely, the introduction of alginate led to a downregulation of TUBB-3 compared against control cells, and the presence of graphene led to a further downregulation (Figure 5 (B)). The alginate control and alginate fiber samples did not have a drastically different TUBB-3 expression, which indicates that the encapsulation process itself did not cause a downregulation of TUBB-3, but that the alginate and graphene-alginate solutions played a larger role. Similarly, the graphene-alginate microfiber samples showed a further downregulation of TUBB-3, which shows that the initial contact with graphene suppressed the health of N27 cells.

Overall, this shows that the introduction of alginate or graphene and the manufacturing process overall will lead to the increase of the dopaminergic activity of N27 cells by upregulating the TH gene, while conversely inhibiting TUBB-3, therefore showing a decreased ability for neurogenesis and axonal maintenance and growth. This is unlike literature values, which show that the seeding of MSCs onto graphene foams increases the presence of the neuronal marker TUBB-3, indicating that MSCs had differentiated into neuronal cells.^[43]^ The inverse finding of the current work indicates that alginate might inhibit the upregulation of TUBB-3 that might occur in the presence of pure graphene.

### 2.7 Genetic Expression of N27 Cells after Prolonged Encapsulation (Day Six)

Cell-laden fibers were cultured for six days to study the long-term genetic effects of encapsulation within alginate and graphene-alginate hydrogels, compared to the genetic effects of the encapsulation process itself. In addition to the previously mentioned TH and TUBB-3, IL-1β and TNF-α were observed in the day six samples. IL-1β is a crucial gene that helps to regulate inflammatory response,^[44]^ and is involved in cell proliferation, differentiation, and apoptosis.^[45]^ Similarly, TNF-α is an inflammatory cytokine that arises during acute inflammation. It can lead to cell death, but it is also an important factor in fighting cancer or infections.^[46]^

As of day six of encapsulation TH levels are still greatly upregulated when compared against the control sample (**Figure 6** (A)); however, these values are lower than seen on day one (Figure 5 (A)). This indicates that the manufacturing process played a greater role in increasing the expression of TH when compared against the long-term encapsulation of cells within either pure alginate or graphene-alginate microfibers. However, in looking at the day six data, one can observe that the alginate controls still exhibit a greater TH expression than the control sample, and that alginate fibers are raised even further. This shows that contact with alginate, and encapsulation within alginate or graphene-alginate microfibers, both increases TH expression, and therefore show an increase in dopaminergic activity. After six days, there is no large difference between the alginate and graphene-alginate fiber samples, indicating that prolonged contact with graphene was not drastically affecting the TH expression of encapsulated cells, and that the degradation of the fiber samples was allowing for greater diffusion into and out of the graphene-alginate microfibers.

**Figure 6.**
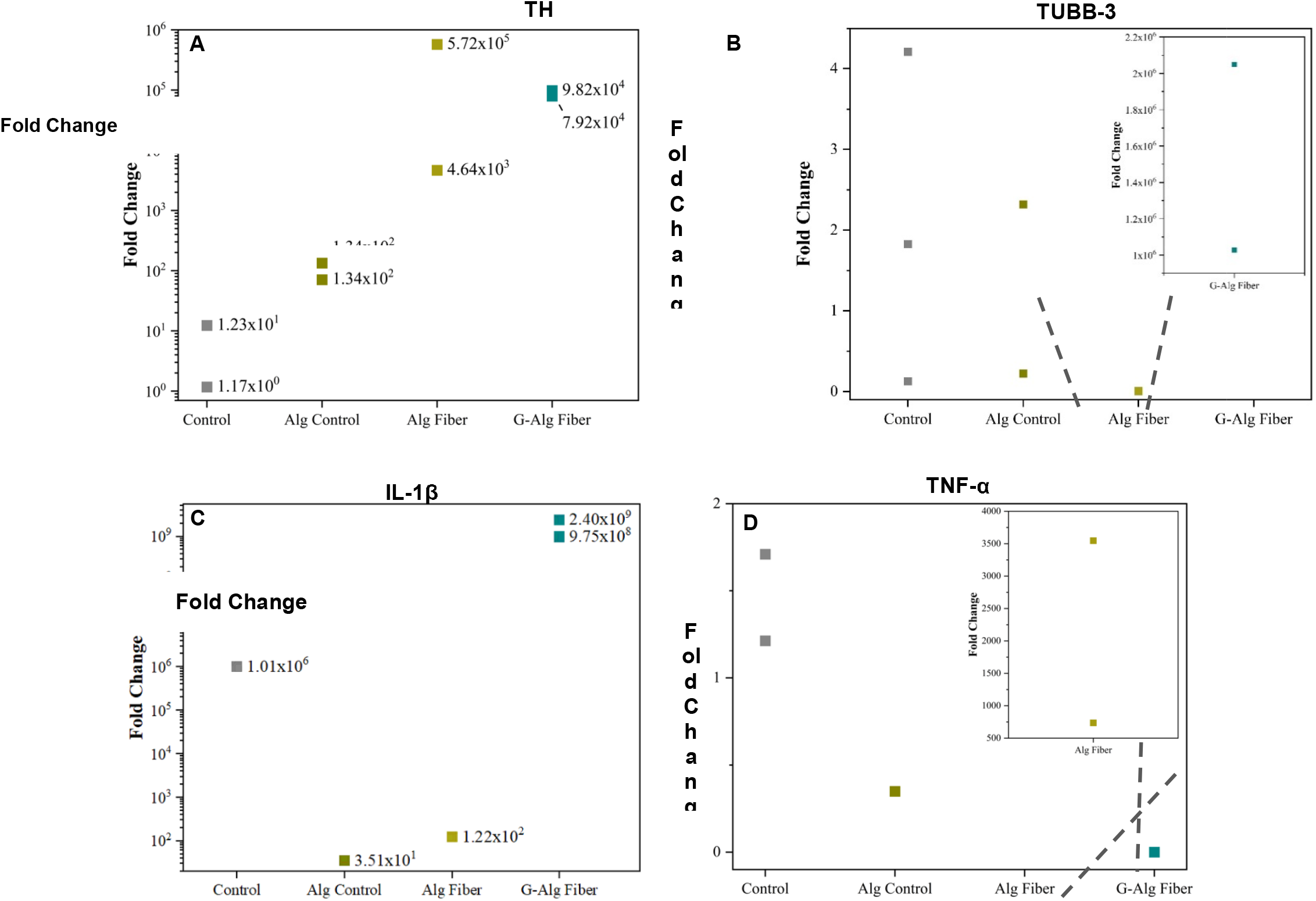
Relative normalized gene expressions (fold changes) within different N27 cell samples after six days, including controls and alginate controls, which were introduced to alginate and stayed within the syringe during the encapsulation procedure before being cultured in a well plate. IL-1β **(A)**, TH **(B)**, TNF-α **(C)**, and TUBB-3 **(D)** levels were analyzed with RT-qPCR (n≥1).

After six days of encapsulation or culturing with alginate, the levels of TUBB-3 are comparable to the levels observed in day one samples, except for the graphene-alginate encapsulated cells, which showed a drastic upregulation of TUBB-3 (Figure 6 (B)). Since this trend was not present in day one, this indicates that while short-lived contact with graphene in graphene-alginate fibers might not cause a change in TUBB-3 expression, long-term contact may lead to a significant change in TUBB-3 levels in N27 cells, which mirrors the findings of Tasnim et al. (2018).^[43]^ Furthermore, this indicates that the actual encapsulation procedure did not cause any extreme changes on TUBB-3 expression levels, but that the greatest parameter to effecting TUBB-3 is prolonged contact with graphene particles within the graphene-laden alginate hydrogels. Since six days in maintenance media leads to some natural degradation in the presence of physiologically relevant levels of sodium, the differences in porosity and diffusion rates between alginate and graphene-alginate microfibers are lessened, but might still affect the TUBB-3 levels after prolonged encapsulation.

As seen in Figure 6 (C), culturing cells with alginate solutions or within alginate microfibers for a prolonged time led to a downregulation of IL-1β when compared against plain cells, as has been observed in literature.^62^ However, encapsulation in graphene-alginate microfibers caused an increase in the expression of IL-1β, indicating that the addition of graphene was triggering an inflammatory response within the cells after six days of co-culture with graphene microfibers. Lack of a similar drastic upregulation within the plain alginate fibers indicates that the interaction between graphene and the cells is the likely cause for the upregulation of IL-1β, as opposed to solely the prolonged entrapment within the alginate hydrogels, or of the decreased porosity or increased size of the graphene-alginate microfibers. Similarly, previous works have shown that graphene-oxide causes caspase-dependent IL-1β expression in THP-knockdown cells, indicative of inflammasome activation, which is relevant to the current findings.^[47]^

After six days of cell culture with the alginate solution, the alginate control cells did not show a very large difference in TNF-α expression, indicating that the presence of alginate alone did not greatly affect the inflammatory response of the cells (Figure 6 (D)). However, after six days of encapsulation within pure alginate microfibers, cells showed drastically higher levels of TNF-α expressions than the alginate controls or the graphene-alginate microfibers, suggesting that perhaps the presence of graphene reduced the inflammatory response of the encapsulated cells. This is an important note; graphene-based sensors are increasingly being used for the sensing and detection of inflammatory cytokines like IL-6, IL-1β, and TNF-α; any concern that graphene might affect the expressions of these genes might skew the resulting data.^[48]^ The present work indicates that graphene might inhibit the expression of TNF-a, but might cause and upregulation of IL-1β, which should be carefully considered before proceeding with graphene-based sensors.

Overall, there are key parameters in the microfluidic fiber fabrication process that lead to different effects on the genetic expressions of N27 cells. For instance, merely introducing the cells into alginate solutions was enough to drastically increase TH levels on day one, a trend which continued through long-term contact with alginate or encapsulation into fibers, and a trend that continued through the six-day observation period, indicating that the dopaminergic activity of encapsulated N27 cells is elevated when compared against the control samples. Similarly, expression levels of TUBB-3 in the alginate control and alginate fibers did not drastically change over the six-day encapsulation, but did greatly increase within the graphene-alginate sample, indicating that the encapsulation process temporarily downregulated TUBB-3 expressions that should have been upregulated in the presence of graphene;^[43]^ furthermore, this effect was temporary, and the graphene-alginate samples showed a large upregulation as of day six. While upregulation of TUBB-3 can indicate the maturity of a neural cell culture, drastic changes in TUBB-3 levels can be problematic, with similar trends appearing in some cancer tissues.^[42]^

## 3. Discussion

This work communicates the successful encapsulation of rat dopaminergic N27 cells into graphene-alginate microfibers using a microfluidic wetspinning technique. Incorporation of graphene into the alginate microfibers increased the conductivity of the microfibers by 148%, leaving the wet hydrogels with a conductivity that is in line with that of native brain tissues. However, further work must be done to accomplish conductivities that will enable the measurements of real-time cell-to-cell communications; increasing the conductivity might be accomplished by adding synthetic polymers, such as PEDOT:PSS.

The incorporation of graphene into alginate solutions led to higher viscosities, which further impacted the physical characteristics of the microfibers by increasing their size and porosities, while decreasing their surface roughnesses. The microfluidic fiber fabrication process showed itself once again to be unmatched for creating reproducible, continuous high-quality microfibers for a variety of applications, including cell encapsulation for the creation of 3D cell scaffoldings for use in tissue modeling and real-time sensing.

The described encapsulation procedure had an encapsulation efficiency rate of 50%, with some cells remaining within the syringe or exciting into the collection bath. Similarly, some cells were able to migrate out of the fiber boundaries, and were observed attached to the bottom of the well plate. The entire six-day observation period showed 30% of cells remaining. This number might be improved on for future works by functionalizing the alginate to further improve the biopolymer for cell adhesion and proliferation.

Before use as a 3D cell construct, it is crucial to understand the genetic implications of encapsulating cells, which prompted the use of RT-qPCR to study the effects of the manufacturing process and long-term encapsulation on the genetic expressions of the genes TH, TUBB-3, IL-1β, and TNF-α. It is important to understand what genetic variations were caused by introduction of alginate, the manufacturing process, and contact with graphene. For this reason, an alginate control, which was mixed and kept within the syringe during encapsulation, was compared against both an alginate fiber sample and a graphene-alginate sample, both on day one and day six.

As of 24 hours after the manufacturing process, the presence of alginate caused a large upregulation of TH levels within N27 cells, with graphene increasing upregulation even further, indicating an increased dopaminergic activity. The opposite trend was true of TUBB-3. However, 144 hours after encapsulation showed a continued upregulation of TH, with the encapsulated samples being more elevated than the alginate controls. TUBB-3 levels were not greatly elevated after six days of encapsulation, but this work shows that the presence with graphene did cause a drastic upregulation in TUBB-3, indicating increased neuronal activity in the presence of graphene in the sample. Similarly, IL-1β was downregulated in the alginate control and pure alginate fiber samples but were greatly elevated for the graphene-alginate encapsulation cells, and TNF-α showed upregulation for the pure alginate fiber samples, indicating that graphene might cause downregulation of that gene, and therefore lead to a decrease in the corresponding signs of inflammation.

The production of cell-laden hydrogels with a conductivity similar to that of native brain tissue can positively influence a number of biomedical applications, such as cellular constructs or scaffolding for regenerative medicines or tissue modeling.^[49]^ However, such systems must be characterized before use, and the effects on cell behavior and genetics must be thoroughly understood. This work aims to elucidate the changes in genetic expressions of N27 cells that have been encapsulated into alginate and graphene-alginate microfibers, and to better understand the full genetic implications of manufacturing cell-laden microfibers, so that they might be successfully utilized for highly complex 3D cell culturing with real-time sensing platforms for analyzing the cellular response of different stimuli.

## 4. Materials and Methods

### 4.1 Manufacturing of PDMS Microfluidic Devices

Devices were created by thermosetting polydimethylsiloxane (PDMS) (Dow Corning, Midland, MI) with photolithographic molds on silicon wafers. Briefly, the microfluidic channel features a main chamber which has dimensions of 130 μm x 390 μm, which holds four chevrons to allow for lateral shaping of the core fluid.^[6, 9, 29, 30, 32]^ These chevrons are 200 μm apart and have dimensions of 130 μm x 100 μm.

PDMS was mixed in a 1:10 ratio of elastomer curing agent to base, and was allowed to sit at room temperature until all air bubbles have risen to the surface. Uncured PDMS was solidified on the molds by cooking it at 80°C for 20 minutes. The two halves were bonded using plasma cleaning (PDC-001, Harrick Plasma, Ithaca, NY, USA) on medium for 20 seconds to activate the surface bonds of the PDMS.

### 4.2 Creation of Alginate and Graphene Solutions

Aqueous graphene solutions were created using established methods.^[9, 22]^ For this study, 4 g of graphite powder (Synthetic graphite powder <20 μm, Aldrich Chemistry, St. Louis, MO) was mixed with 200 mL of DI water, along with 400 mg of BSA (A7906, Sigma-Aldrich, St. Louis, MO), and the resulting mixture was placed into a standard kitchen blender for 45 minutes in 10 minute intervals to prevent overheating. 3.5% (g/mL) alginate (very low viscosity, Alfa Aesar, Ward Hill, MA, USA) was dissolved into either DI water or graphene for alginate or graphene-alginate solutions, respectively.

### 4.3 Viscosity Measurements

To measure the viscosity, 10 mL of both pure 3.5% alginate and 3.5% alginate and the graphene solution were mixed at 40 °C for a minimum of one hour. A Canon-Fenske viscometer (350 727H, Canon Instrument Company, State College, PA, USA) was suspended in a water bath maintained at 40 °C to ensure constant temperature. The solutions were loaded into the viscometer and were allowed to rest for 30 minutes to rise to 40 °C, at which point they were introduced into the efflux bulb via suction. The time required for the liquid line to fall between the starting point and the ending point (the efflux time) was measured, and the viscosity was calculated with the following equation, where *v* is the kinematic viscosity, *C* is the viscometer constant, and *t* is the efflux time.

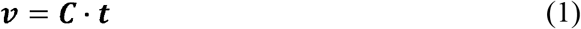

### 4.4 Fiber Creation

To create the microfibers, both the core (3.5% alginate in DIW or graphene) and sheath (25% PEG and 0.08% CaCl_2_·2H_2_O in DIW) solutions were placed into syringes and were injected into the microfluidic device using a double syringe pump (Cole Parmer, Vernon Hills, IL, USA). The sheath solution was introduced into its two inlets using a Y-shaped tubing splitter, and the flow rate ratio was 200:20 μL min^−1^ (Sheath : Core). Fibers were collected from a bath of 15% CaCl_2_·2H_2_O in DIW.

### 4.5 SEM Characterization

A JEOL JCM-6000 Benchtop Scanning Electron Microscope (JEOL, Akishima, Tokyo, Japan) was used to view the cross-sectional and longitudinal topographies of the microfibers. Fibers were dried at room temperature (RT) overnight and were mounted onto electrically conductive electrical tape (Electron Microscopy Services, Hartford, PA, USA).

### 4.6 Profilometry and Surface Roughness

A NewView 7100 Profilometer (Zygo, Middlefield, CT, USA) was used to quantify the surface topology of alginate and graphene-alginate microfibers. Once dried at RT overnight, fibers were placed onto a glass coverslip and were observed with the Zygo MetroPro system to generate the average roughness value, or the R_a_ value.

### 4.7 Porosity Calculations

Fibers were fabricated and dried overnight at RT. Glass cover slides were weighed before the dry weight of the microfibers was gathered. Fibers were soaked in DI water (DIW) overnight, and the next day, excess water was removed using a Kimtech wipe (Kimberly-Clark Global Sales, LLC, Roswell, GA, USA), and the wet weight was measured. The porosity was calculated using the following equation, where *M_w_* is the wet weight, *M_d_* is the dry weight, *ρ* is the density of the soaking liquid, and *V* is the volume of the wet fibers.

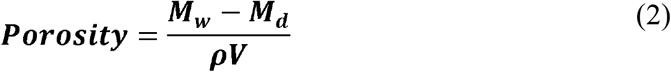

### 4.8 Electrical Characterizations

Fibers were fabricated in a non-sterile environment and were allowed to dry overnight at RT on paper frames. Once dry, between 20 and 30 fibers were mounted onto plastic cover slips using two strips of double-sided conductively adhesive carbon NEM tape (Nisshin Em. CO., LTD., Mitaka, Tokyo, Japan) with a 1-2 mm gap. Fibers were further ohmically connected using a conductive silver paste (PELCO^®^ Conductive Silver 187, Ted Pella, Inc., Redding, CA, USA).

A VersaSTAT 4 Potentiostat (Princeton Applied Research, Oak Ridge, TN, USA) was used to conduct cyclic voltammetry on mounted alginate and graphene-alginate microfibers, thereby yielding a current vs. potential graph that allowed for the calculation of the resistance of the fibers. From there, the conductivity of the fibers was calculated with the following equation, where Ω is the conductivity, *R* is the resistance of the fibers, *N* is the number of mounted fibers, and *A* is the cross-sectional area of a single fiber.

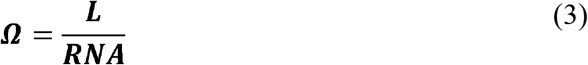

### 4.9 Cell Culturing Procedures

N27 cells with a passage number of less than 10 were used for this study. They were cultured using standard techniques at 37 °C in a 5% CO_2_ atmosphere, and were kept in maintenance media (MM) created with RPMI Medium 1640 (1X) (Gibco Life Technologies Ltd., Paisley, UK), which was supplemented with 10% fetal bovine serum (FBS) (One Shot format, ThermoFisher Scientific, Waltham, MA, USA), 1% penicillin (10,000 U mL^−1^)-streptomycin (10,000 μg mL^−1^) (Gibco, Waltham, MA, USA), and 1% L-glutamine 200 mM (100X) (Gibco Life Technologies Corporation, Grand Island, NY, USA). Cells were passaged at 70% confluency.

### 4.10 Cell Encapsulation

Alginate powder was sterilized with 70% ethanol, and 0.07g was dissolved into 1.4 mL of either sterile DIW or sterile graphene solution at 40 °C with a heated magnetic stirrer for a minimum of one hour until homogeneous. Cells were trypsinized, and 0.6 mL of cell suspension with a concentration of 6 x 10^6^ cells mL^−1^ was added to form 2 mL of a cell-alginate or cell-alginate-graphene solution with a final cell density of 2 x 10^6^ cell mL^−1^.

Fiber fabrication proceeded as described above, this time in sterile conditions under a biological fume hood. Both the sheath and bath solutions were autoclaved, and microfluidic devices were flushed with 70% ethanol to sterilize them before use.

### 4.11 Live-Dead Cell Assay

To calculate the amount of live cells within the fibers, first, fibers were removed and were placed into a 1.5 mL centrifuge tube. All excess media was removed, and the fibers were weighed. Fibers were dissolved with 0.1 M phosphate buffered solution (PBS) by placing 0.5 mL of media and 0.5 mL of PBS and pipetting repeatedly. If needed, dissolving fibers were placed into the incubator for up to 30 minutes to allow for degradation.

Once fibers were dissolved, the centrifuge tubes were centrifuged for five minutes according to the manufacturer protocol. Excess PBS/MM was removed, and the remaining pellet was suspended into fresh MM. A trypan blue exclusion allowed for the measurement of the cells within the suspension, which allowed for calculation of the concentration of cells per gram of fiber.

### 4.12 RNA Isolation and cDNA Synthesis

Samples were frozen at the desired timepoints and were stored at −20 °C until RNA isolation was performed. To isolate RNA, the TRIZOL™ Plus RNA Purification Kit from Ambion was used according to the manufacturer protocol (ThermoFisher Scientific, Waltham, MA, USA). RNAseZap™ from Invitrogen™ (ThermoFisher Scientific) was used to thoroughly clean all surfaces and tools before beginning. Isolated RNA was checked with a NanoDrop and the concentration of all samples was normalized before continuing.

Invitrogen™ Superscript™ IV VILO™ Master Mix with ezDNAse™ enzyme (ThermoFisher Scientific, Waltham, MA, USA) was used according to manufacturer protocol to convert RNA into cDNA.

### 4.13 RT-qPCR Analysis

Real-time RT-qPCR was carried out using Taqman™ assays from ThermoFisher Scientific (Waltham, MA, USA). The following probes were used for each gene: B2M (Cat #: 4453320, Assay ID: Rn00560865), IL-1β (Cat #: 4453320, Assay ID: RN00580432_m1), IL-6 (Cat #: 4453320, Assay ID: RN01410330_m1), TH (Cat #: 4453320, Assay ID: RN00562500_m1), TNF-α (Cat #: 4453320, Assay ID: RN99999017_m1), and TUBB-3 (Cat #: 4448892, Assay ID: Rn01431594_m1). The Taqman™ Fast Advanced Master Mix (ThermoFisher Scientific) was used according to manufacturer protocol. Eighty standard curve reactions were used according to the QuantStudio™ Design&Analyze Software (v1.5.1, ThermoFisher Scientific) and a QuantStudio™ 3 Real-Time PCR System (ThermoFisher Scientific).

### 4.14 Statistical analysis

Analyses were carried out using R Statistical Software (v 4.0.3) was used in conjunction with RStudio Desktop (v 1.3.1093) to carry out Analysis of Variance (ANOVA) tests and Tukey tests to compare the means across samples.

## Acknowledgments

This work was partially supported by the Office of Naval Research (ONR) Grant N000141612246, ONR Grant N000141712620, and National Science Foundation Grant 2014346. We thank Dr. Farrokh Sharifi, Kelli R. Williams, and Dr. Jie Luo for their assistance, and Dr. Anumantha Kanthasamy for the gift of rat dopaminergic neural cells (N27s).

## Author Contributions

M.C.M. and A.E.N.A. carried out the experimental studies in this work. M.C.M. processed the experimental data. R.L.P. and A.H.W. were involved in design of the experiments and consulted regarding the manuscript. R.M. and N.N.H. conceived the study, acquired funding, and supervised the work. M.C.M., R.M. and N.N.H. wrote the manuscript.

## Competing Interests

The authors declare no competing interests for this work.

## Conductive Cellular Scaffolding

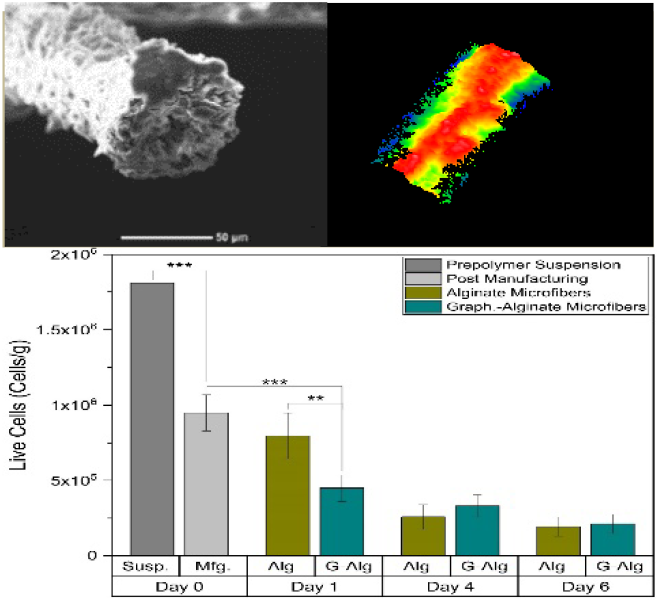
Graphene is used to increase the conductivity of biocompatible alginate microfibers for use in encapsulating dopaminergic rat N27 cells. The conductivity of the microfibers is increased by 143%. The genetic effects of encapsulation are studied to better understand the use of hydrogel scaffoldings as cell constructs.

## Notes

### Competing Interest Statement

The authors have declared no competing interest.

